# Integrative analysis of fine-scale local adaptation of winter moths to variable oak phenology

**DOI:** 10.1101/2025.10.10.681598

**Authors:** Rona Learmonth, Andrea Estandía, Léa Beaupère, Ella F. Cole, Ben C. Sheldon

**Affiliations:** Edward Grey Institute of Field Ornithology, Department of Biology, University of Oxford, United Kingdom

**Keywords:** adaptation, synchrony, phenology, moth, Lepidoptera, carry-over effects

## Abstract

For herbivorous insects whose fitness depends on tight phenological synchrony with host plants, spatial variation in plant phenology can impose strong selective pressures and promote local adaptation to host timing. These dynamics are central to predicting how species will respond to environmental change, particularly climate-driven shifts in plant phenology. The winter moth (*Operophtera brumata*) relies on synchronising larval egg hatch with leaf budburst of deciduous trees, yet whether they are locally adapted to their hosts’ phenology, and their capacity to track future change, remains unclear. Here, we investigated potential small-scale local adaptation of winter moths to oak tree phenology in Wytham Woods, UK, a 385-hectare woodland, within which oak budburst can vary by up to three weeks within a given year. We conducted laboratory temperature manipulation experiments using 76 clutches across six temperature treatments, and field translocation experiments using over 200 clutches. We combined these experiments with assessment of population structure from whole-genome sequencing of 59 individuals. This integrative approach allowed us to assess local adaptation in terms of phenotypic differences, fitness consequences, and genetic evidence. Temperature manipulations revealed systematic differences in the timing of egg hatching across temperature treatments at the clutch level which were linked to carry-over effects from the mother’s emergence time, but unrelated to their source tree budburst timing. Field translocation experiments further showed no significant differences in survival of individuals transplanted to trees with phenology differing from their original host tree, and there was no genetic structure across the population. Together, these results reveal consistent differences in hatching phenology despite the absence of population structure, strong selection, or accordance with relative tree phenology. Our findings advance our understanding of the mechanisms maintaining close synchrony in trophic interactions at small scales, which may drive spatial variation in evolutionary responses to future climate change.

## Introduction

Phenology, the timing of life-cycle events such as flowering in plants or hatching in animals, determines species interactions [1]. When a species’ survival depends on matching its timing with another, such as a consumer relying on an ephemeral food resource, there is strong selection for synchrony [2,3]. This is especially important for taxa such as phytophagous insects which often associate with a single host plant across their lifespan. If they fail to synchronise with leaf production of their host plant, their fitness and survival can suffer [4]. Because of this, phytophagous insects face strong selection to match their own life cycle timing with the timing of the plant’s growth [5–7].

Plant phenology exhibits substantial spatial variation driven by differences in temperature, precipitation, photoperiod, and elevation, as well as genetics [8]. These differences create a mosaic of phenological environments across landscapes, with spring leaf emergence sometimes varying by weeks or even months across relatively short distances [9]. Additionally, individual host phenology is very consistent across years [9], creating predictable conditions for insect populations that must synchronise their life cycles with local hosts. Synchrony can be achieved through multiple mechanisms, including plasticity in developmental responses, maternal effects, and local adaptation [10,11]. Distinguishing among these pathways is critical: while plastic shifts may allow adjustments to shifting host timing in the short term, populations with high genetic variation and moderate local adaptation may indicate greater ability to rapidly undergo genetic adaptation to match directional shifts in host phenology [12]. In extreme cases, such local adaptation can even lead to the formation of new species [13].

Investigating the mechanisms of adaptation to phenology is important in the context of climate change. Rising temperatures are driving advances in phenology across taxa such as birds [14,15] and insects [16]. These shifts can disrupt synchrony when interacting species differ in their environmental cues or degrees of plasticity [17]. Understanding the mechanisms by which phytophagous insects maintain synchrony with their local hosts across space can help us predict how they may respond to climate-mediated shifts through time [18].

Most evidence for local adaptation comes from large spatial scales, such as across latitudinal gradients [7]. For example, the arctiid moth (*Utetheisa ornatrix*) showed local adaptation to its host plant at continental but not regional scales [19]. Different factors may drive local adaptation at different spatial scales, including complex interactions between the species’ dispersal, the strength of selection, environmental heterogeneity, and demographic factors such as population size fluctuations [20,21]. Thus, understanding the scales at which local adaptation arises is a key step in untangling the processes underlying synchrony in phenologically variable landscapes.

The adaptive deme formation (ADF) hypothesis provides a mechanism for small-scale local adaptation within host species. ADF proposes that phytophagous insects can adapt to individual host plants, forming locally adapted ‘demes’—spatially discrete populations with random internal breeding [22–24]. Originally, ADF was attributed to adaptation to individual host plant defences [23], but this received weak empirical support [24]. More recently, intraspecific variation in host phenology has been proposed as an ADF driver [6]. Komatsu & Akimoto (1995) provided robust evidence for phenology-mediated ADF in the aphid (*Kaltenbachiella japonica*), demonstrating local adaptation to budburst timing on individual host trees [5]. It has also been suggested that synchrony between winter moths (*Operophtera brumata*) and their host oak (*Quercus robur*) trees is maintained by ADF [25].

Winter moths are ideal for studying local adaptation to vegetation phenology. They have demonstrated rapid climate-mediated genetic evolution by advancing their phenology to match that of their host trees over just one decade of warming [26,27]. This capacity for rapid genetic adaptation highlights the importance of understanding the mechanisms underlying their phenological synchrony. Synchronising egg hatching with host tree budburst is critical for larval survival: hatching too early leads to starvation due to the absence of suitable leaves, whereas hatching too late results in feeding on leaves that have already accumulated defensive compounds such as tannins and become tougher and less nutritious, reducing survival and fitness [28,29]. If winter moths can maintain synchrony through fine-scale local adaptation, this could make them resilient to future climate change by sustaining variation in hatching times within populations, which can buffer against spatially heterogeneous climate impacts. The female adults are brachypterous, with very short wings incapable of flight, which is likely to limit dispersal and gene flow, creating conditions favourable for local adaptation [25]. Indeed, studies have found strong population genetic structure across Europe [30,31], but these were conducted at relatively coarse spatial scales, leaving a gap in our understanding of whether fine-scale genetic adaptation can occur to individual hosts.

Wytham Woods, a 385-hectare mixed deciduous woodland in Oxfordshire, UK, is an appropriate site to test for smallscale local adaptation in winter moths, combining decades of ecological research [32,33] with fine-scale variation in oak budburst timing, which can vary by over 20 days across the woodland [9]. This microhabitat variation creates a natural setup to test small-scale adaptation to host phenology.

Here, we test whether winter moths show small-scale local adaptation to the phenology of individual host trees. Specifically, we ask whether differences in oak budburst timing correspond to adaptive genetic differences in moth hatching phenology. We address this by combining three complementary approaches: (i) common garden experiments under controlled temperatures to test for phenotypic differences, (ii) translocation experiments to measure fitness consequences of phenological mismatch, and (iii) genomic analyses to evaluate population structure and the potential for fine-scale adaptation. This combination of methods provides a novel test of small-scale phenological adaptation in phytophagous insects.

## Methods

### Temperature manipulation experiment

We conducted a temperature manipulation experiment using common gardens at six temperature conditions to assess local adaptation. If local adaptation was present, we predicted that clutches from different trees would show consistent patterns of phenology linked to the timing of their source tree due to underlying genetic differences when in a common garden environment. For example, a clutch from an early tree would consistently hatch earlier relative to clutches from other trees across multiple temperature conditions, independent of temperature-mediated phenotypic plasticity.

#### Field sampling

Individuals used in this experiment were the eggs of females caught from Wytham Woods, Oxford, in November and December 2024. Females were caught in paired lobster pot traps, which capture emerging winter moths as they climb up the tree trunk and are widely used for geometrid moth sampling [34]. These traps were installed on the north and south-facing sides of 10 pairs of oaks (*Quercus sp*.) trees along the main path in the woods (Fig. 2a). These 20 trees effectively made up a transect across the woods, supplemented by four additional trees included as part of long-term monitoring. Trees were selected to cover a normal distribution of phenology, with a 29-day range in their 2024 half-dates (Fig. S1), used as a proxy for budburst. Half-dates were derived from time-series of multispectral data collected using drones, which were flown over Wytham Woods every three days, weather permitting, from March to May each year (see [35] for full details). The NDVI images from these flights were stitched into orthomosaics, from which tree crowns for the focal oaks were manually delineated. For each tree, mean daily NDVI values for the central 75% of the tree crown were extracted, and the date at which NDVI reached 50% in the green-up curve identified as a proxy for budburst date [36]. Each trap was checked three times per week, removing all organisms from the trap and collecting only the winter moths present. Females were housed individually in falcon tubes with a roll of tissue for oviposition and stored outside at ambient conditions. The eggs were maintained outside at ambient conditions until January, when they were transferred to treatment groups. All clutches with a minimum of 30 eggs were used in the common garden experiment (n=76), with at least six clutches present per tree. To relate timing of subsequent egg hatching to source tree phenology, we monitored the timing of budburst at the source trees in spring 2025, when the eggs would hatch. We conducted budburst scoring at three-day resolution following the seven-step key for oak bud development created by [37] as used by [9].

**Fig. 1.**
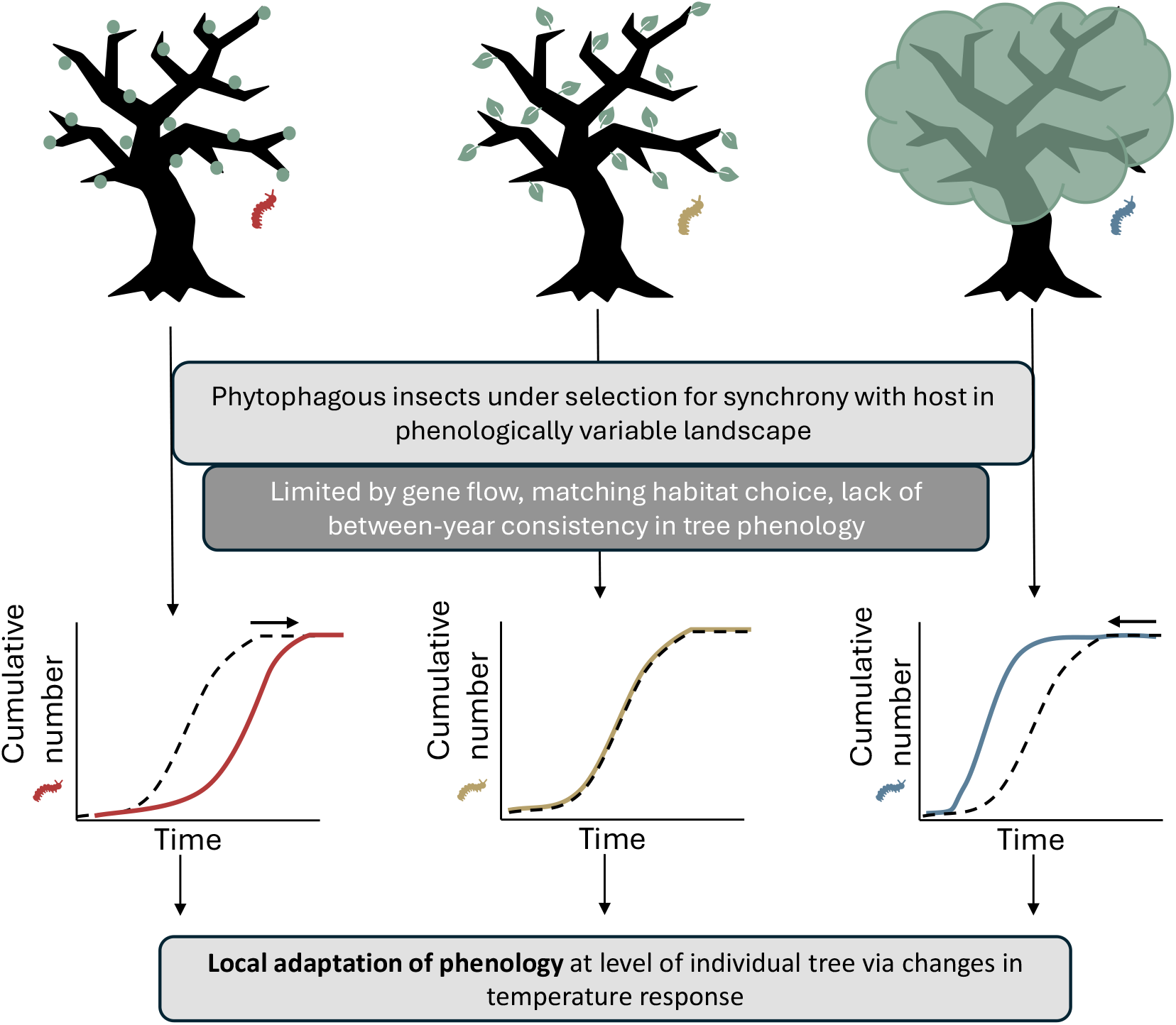
Conceptual illustration of how variation in plant phenology may drive adaptive deme formation, or local adaptation at the level of individual host plants. The three trees indicate a phenologically variable landscape, where they differ in their spring budburst phenology. The tree on the left is relatively late, with buds at an early stage of development, while the middle tree is in budburst, and the early tree on the right is already fully in leaf. If the insects on each of these trees emerged simultaneously, matching the optimum hatching phenotype for the average budburst phenology of the woodland, the red and blue caterpillars would be asynchronous with their own host tree. The red insect would starve on its tree, while the blue insect would face unpalatable leaves with high tannin content. If caterpillars that emerge on their host tree too early or too late cannot undertake costly dispersal to another tree which might better match their phenology, they may starve. Thus, we can expect strong selection for the phytophagous insects to match the timing of their host trees if trees have between-year consistency in their relative phenology. This selection is expected to result in a shift in the temperature-mediated phenology, such that clutches from early trees hatch earlier under shared environmental conditions. This is shown on the three hatch curves, where the dotted line indicates the mean expected hatching response under shared conditions, while the coloured line in each case shows how we expect this response to be modified at each tree due to selection for synchrony, further indicated by the direction of the black arrow.

**Fig. 2.**
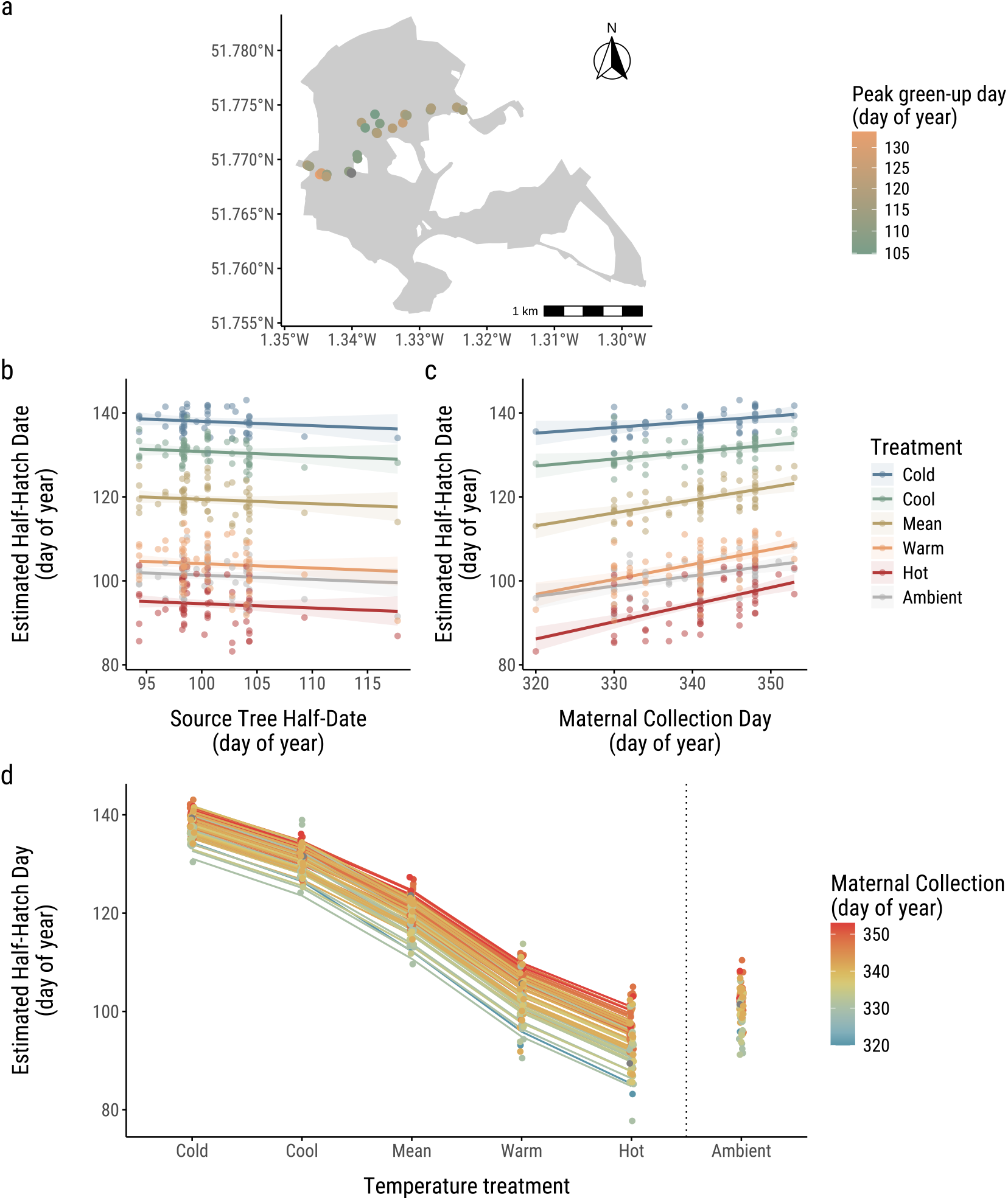
Results of temperature manipulation experiment. Panel (a) shows a map of trees sampled for adult winter moths in Wytham Woods, Oxford. The woods are shown in lighter grey, with the paths marked out by darker grey lines. Dots indicate trees sampled, where the colour indicates the relative budburst of the tree in 2024 based on the date of maximum increase in NDVI. Panel (b) shows the conditional effects estimate of the effect of source tree budburst day on egg half-hatch day, both shown as day of year. Individual coloured lines represent this relationship at each experimental temperature, surrounded by the shaded area of the 95% confidence intervals. Panel (c) similarly shows the conditional effects estimate of the link between maternal collection day and egg half-hatch day. As previously, the coloured lines indicate the relationship at each experimental temperature, surrounded by 95% confidence intervals. In each case, the conditional effects plot represents the relationship between the axes when all other factors included in the model are controlled. Each treatment contains one replicate at each temperature of each of 100 clutches. Lastly, panel (d) shows a conditional effects reaction norm of estimated half-hatch timing of clutches across six temperature treatments. The figure shows the raw half-hatch date of individual clutches, represented by circles, at each of the six temperature treatments (cold (-2.5 SD), cool (-1.5 SD), mean, warm (+1.5 SD), hot (+2.5 SD), and ambient (outdoor)). The lines represent the reaction norm for how estimated half-hatch day changes with temperature for each clutch (n=76). In each case, the line represents the conditional effect when all other factors included in the model (source tree budburst, maternal collection day, interactions of collection day with treatment) are controlled. The ambient treatment differed from all others in that clutches experienced full natural temperature variation, rather than the matching daily regimes experienced by all experimental treatments, and is therefore not connected in the reaction norm. The lines and points are coloured on a continuous scale by maternal collection day.

#### Experimental set-up

The temperature manipulation set-up used five incubators (PHCbi MLR-352-PE), which represented treatment groups cold, cool, mean, warm, and hot. The temperature data used to generate these treatments was taken from the daily maximum, minimum, and mean HadUK-Grid Gridded Climate Observations on a 5km grid v1.3.0.ceda (1965-2016) [38] for the 5km grid square covering Wytham Woods. We used the function `stack_hourly_temps` in the *chillr* package [39] to interpolate hourly temperatures from the daily maximum and minimum, then averaged these across years. Due to the limitations of the incubator programme memory, we averaged across each two-hour increment, which were each averaged across two-day periods. This resulted in 24-hour 12-step programmes, each repeated for two cycles representing the mean treatment. The other temperature treatments were then calculated by adding or subtracting standard deviations of monthly mean temperature across 50 years as a constant. This resulted in the hot (+2.5 SD), warm (+1.5 SD), cool (-1.5 SD), and cold (-2.5 SD) treatments (see Fig. S2). Lighting was equal across all chambers, set to the nearest monthly mean photoperiod in terms of lighting being on or off for each two-hour period. We also included a sixth ambient treatment, where sub-clutches were kept in a clear box outside in a shaded area and experienced natural temperature variations. We monitored egg hatching weekly from mid-February, then twice weekly from the beginning of March. Egg colour changes from orange to black two to three days before hatching. Egg hatch checks were switched to daily from the first egg darkening, and continued until all eggs were hatched. At each check, the cumulative number of hatched larvae in each sub-clutch was counted. All larvae were reared in situ for use in other experiments.

#### Data analysis

We conducted all data analyses in R [40]. For the temperature manipulation experiment, we first calculated the proportion of larvae hatched over time. To account for death of some larvae and hatching of others, we recalculated the cumulative total as the sum of the positive daily differences. The daily proportion hatched was then taken as that day’s cumulative total divided by the maximum cumulative total, rather than the highest day’s count. This method corrected for deaths during development and resulted in a total percentage hatched of 90.7%, similar to the 90.0% calculated in a previous year [41], and the high estimates seen in other experiments [4]. We then produced logistic hatch curves of the proportion of eggs hatched by experimental day for each sub-clutch in each treatment using the R nonlinear least squares (`nls`) function in the *stats* package [40]. From these curves, we extracted the half-hatch day at which 50% of eggs had hatched, a measure of mean hatch phenology for a sub-clutch. Using the *brms* package [42], we modelled how half-hatch day was impacted by temperature treatment, maternal collection date, and 2025 tree budburst date, as well as interactions of temperature treatment with collection date and budburst. Maternal collection date was taken as a proxy for the mother’s timing of adult emergence to investigate potential maternal carry-over effects in phenology. Using `LOO-CV` in the *brms* package, the best model was selected as one with all variables except the interaction of budburst date with temperature treatment. The model included female identity as a group-level effect, and allowed the intercept of treatment to vary among females. We tested models which also allowed the slope of treatment to vary among females but as each female was represented only once per treatment, there was insufficient within-group data to estimate slope variance. Therefore, here we present only the random intercept model. Each model was run using a Gaussian distribution on four Markov chains, with 5000 iterations each. Posterior predictive checks comparing 1000 draws of the posterior predictive distribution to the distribution of observed data showed strong agreement, indicating the model reflects real patterns in the data.

### Translocations

#### Field sampling

We conducted a translocation experiment to assess evidence for local adaptation to individual trees through fitness differences after translocation. Eggs for this experiment were the offspring of females sampled from November-December 2023 in Wytham Woods using paired lobster pot traps on the north and south-facing sides of 26 oaks (*Quercus robur*) spaced 5-238 m apart, covering a 2.4 ha area (Fig. 3a). We chose these trees as their budburst and winter moth larval phenology had been recorded in the preceding spring and were known to span a range of dates. We checked traps three times per week, transferring all females to individual tubes with strips of cardboard and tissue paper as egg laying substrates. If available, a male caught in the same trap on the same day was placed with the female to ensure mating. Females were stored in an outside shed at ambient conditions for mating. After three weeks, we removed dead adults and stored the eggs outside at natural temperatures until late February. We collected a total of 1112 winter moths, of which 218 females were allowed to oviposit, resulting in 169 clutches of viable eggs. Of females allowed to oviposit without a male, 70/95 produced viable eggs, compared to 99/123 of those stored with males, suggesting that the majority of females had mated prior to capture.

**Fig. 3.**
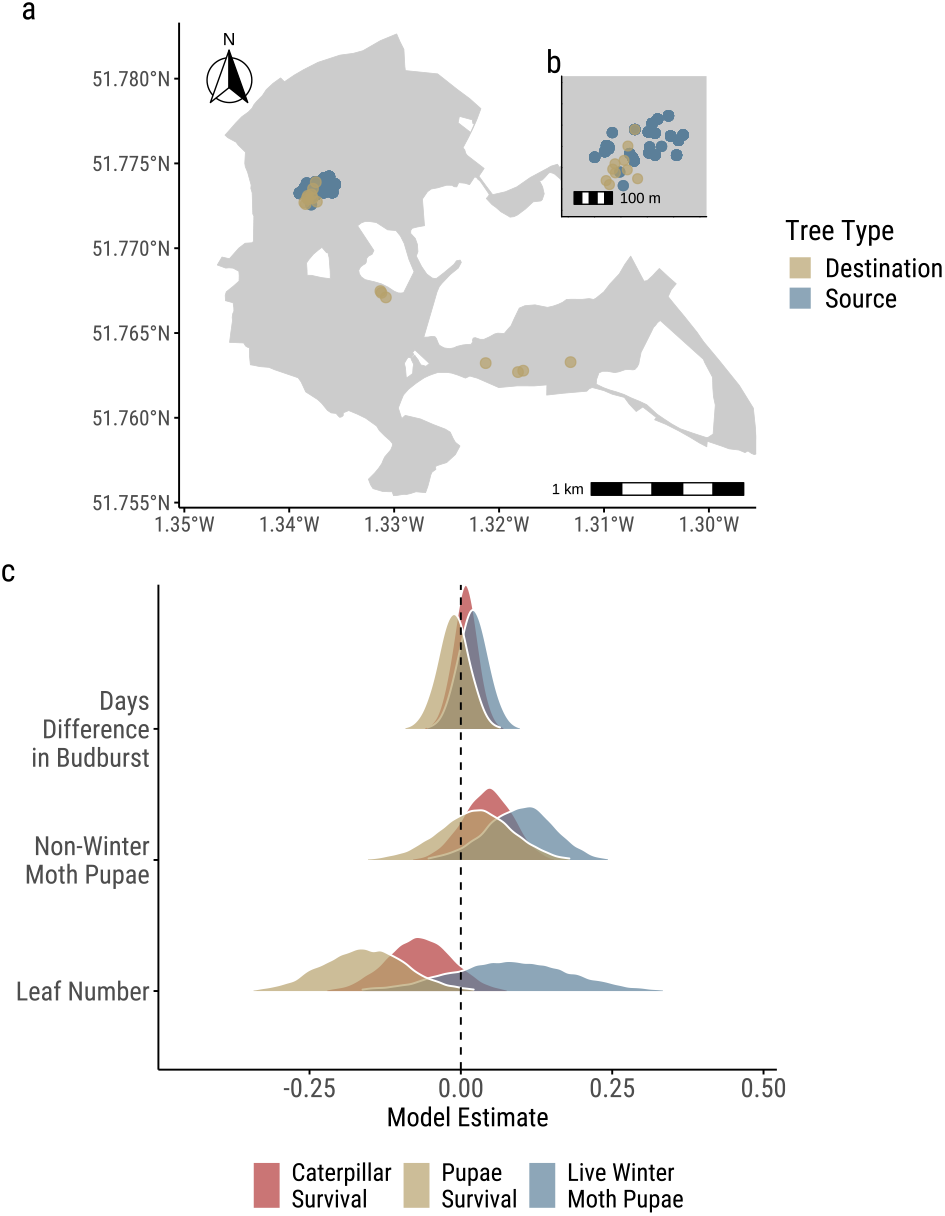
Translocation experiment results. Panel (a) shows the map of where clutches of moths were translocated from source (blue) to destination (yellow) trees within Wytham Woods, with an inset (b) showing a zoomed-in section of the woods which was more heavily sampled. Panel (c) shows the conditional effects ridge plot of posterior distributions for translocated winter moth egg survival to caterpillar stage (red), pupae stage yellow), and live pupae at the end of the experiment (blue). For days difference in budburst, the distribution in each case is closely centred around 0, indicating no significant fitness effects. Posterior distributions were extracted from model estimates when all other factors were controlled. The effects of translocation on pupal mass are narrowly distributed around zero, and are therefore not shown to avoid distorting the plot.

#### Experimental set-up

We translocated each clutch (n=122) from the tree of maternal capture to a matched and mismatched treatment, see Figure 3a. Translocation trees were selected based on (i) their accessibility (within 5m of a path) to the 21m high mobile elevating working platform (MEWP) used to deploy the winter moths, and (ii) their phenology based on NDVI scores derived from drone footage, as previously. Clutches from each tree were assigned to ‘match’ and ‘mismatch’ treatments. The source parental tree was taken as the ‘match’ treatment wherever possible, with an alternative tree which minimised the difference in peak green-up used if the source tree was inaccessible. Mismatch trees were selected from those accessible trees which maximised the positive (‘late’) or negative (‘early’) differences in peak green-up from the source tree. The mean difference in peak green-up was 0.53 (± 0.29 SE) days for match trees, -5.02 (± 0.76 SE) for early, and 4.56 (± 0.49 SE) for late. As trees show consistent between-year differences in their phenology [9], these treatments selected based on 2023 data were assumed to correspond to the relative phenology the eggs would experience in 2024. The actual budburst phenology experienced by the larvae in 2024 was collected by ground-based bud scoring (see [9] for full methods) and used in subsequent analyses. Tree phenology derived from NDVI green-up curves and budburst scoring have been shown to be comparable (Morley et al., 2025). In early March, we deployed sub-clutches of 12 eggs in Eppendorf tubes within mesh bags (200-micron, 20 x 30 cm) around tree branches at similar heights (approx. 10m) within each translocation tree in the densest section of middle-upper canopy reachable from the MEWP. We secured bags with plastic cable ties before opening Eppendorf tubes within the sealed bags to prevent any eggs being lost. We also secured 22 control bags, which contained no winter moth eggs, with at least one bag per tree. These were used to assess the background population density of winter moths on the tree, and so to indicate whether winter moths recovered from the bags at the end of the experiment were likely those we artificially introduced. Across the 22 control bags, the only winter moth present was one dead pupa, indicating that few, if any, non-experimental winter moths were present on branches. This may imply that the introduced populations experienced higher population density than is typical or that bags were placed outside the areas on which eggs tend to be laid, or that they prevented immigration of caterpillars hatched elsewhere. In early June, we used the MEWP to collect the bagged branches and counted the number of live or dead winter moth caterpillars and pupae to assess survival of the introduced eggs through life stages. We also weighed all live winter moth pupae recovered as an accepted proxy for expected female fecundity [4]. In addition, we identified and recorded the numbers of live and dead non-winter moth caterpillars and pupae to assess potential levels of competition. To account for differences in resource abundance between bags, we counted the total number of leaves and made mean herbivory estimates across three randomly selected leaves from each bag using the LeafByte app [43]. From a photo of a leaf taken against a white background with marked 10cm scale, this app removes background from images, then calculates leaf area and percentage herbivory. The final sample size comprised 222 experimental bags and 21 control bags across 17 translocation trees, representing the clutches of 119 individual females from 23 source trees. A total of 22 bags (15.7%) were lost or excluded from subsequent analyses due to high winds, failure to pop the Eppendorf in the bag, or branches snapping before budburst.

#### Data analysis

We modelled how winter moth fitness was affected by differences in budburst phenology compared to the home tree to assess local adaptation in terms of local vs foreign and home vs away criteria [44]. We used *brms* to model the impact of difference in budburst date on four proxies for fitness: (a) the number of individuals surviving to caterpillar stage, estimated as the number of dead and live caterpillars and pupae present; (b) the number of individuals surviving to the pupal stage, made up of the dead and live pupae present; the number of live pupae at the end of the experiment, and (d) the mean weight of live pupae. See Figure S1 for correlations among these fitness estimates. We investigated separately how each fitness metric was affected by the days difference in 2024 budburst at their translocation tree relative to their maternal source tree as a linear and quadratic term to account for potential differences in the effect of being early vs late; the z-scaled number of non-winter moth Lepidoptera present, and number of leaves present in the translocation bag were used to account for interspecific competition and resource availability respectively. Transplant and source tree identities were included as group-level effects. The models for survival numbers used zero-inflated Poisson distributions, while the weight model used a Gaussian distribution. All models were run on four Markov chains, each with 5000 iterations. We used leave-one-out cross-validation (LOO-CV) for model comparison to select the best model for each fitness component. We used posterior predictive checks to assess the model’s fit to the observed data, with a draw of 1000 replicated datasets from the model’s posterior distribution showing good visual agreement with the observed data in density plots. In addition, to assess if artificial introduction of winter moth caterpillars impacted food availability, we used a simple linear model to test whether percentage herbivory differed systematically between experimental and control bags.

### Genomic analyses

#### Genomic sample collection

We collected 59 winter moth caterpillars using water traps and cutting branches from oak trees (n=8) in Wytham Woods, during the spring of 2023 (Fig. 4a). We focused on six oaks, organised into three pairs, with each pair consisting of two trees located within 50 metres of each other and varying in budburst days by at least 15 days. This spatial arrangement allowed us to explore: (i) population structure and gene flow, the effects of distance for winter moth populations inhabiting trees with similar phenology but distant in space, the effects of phenological variability for winter moth populations from trees close in space but with differing phenologies. We also sampled two additional oaks to represent the eastern section of the woodland and explore any potential isolation by distance pattern. We collected between three and ten caterpillars from each tree, ensuring representation across all eight trees. Immediately after collection, we preserved the caterpillars at -80°C in individual Eppendorf tubes.

**Fig. 4.**
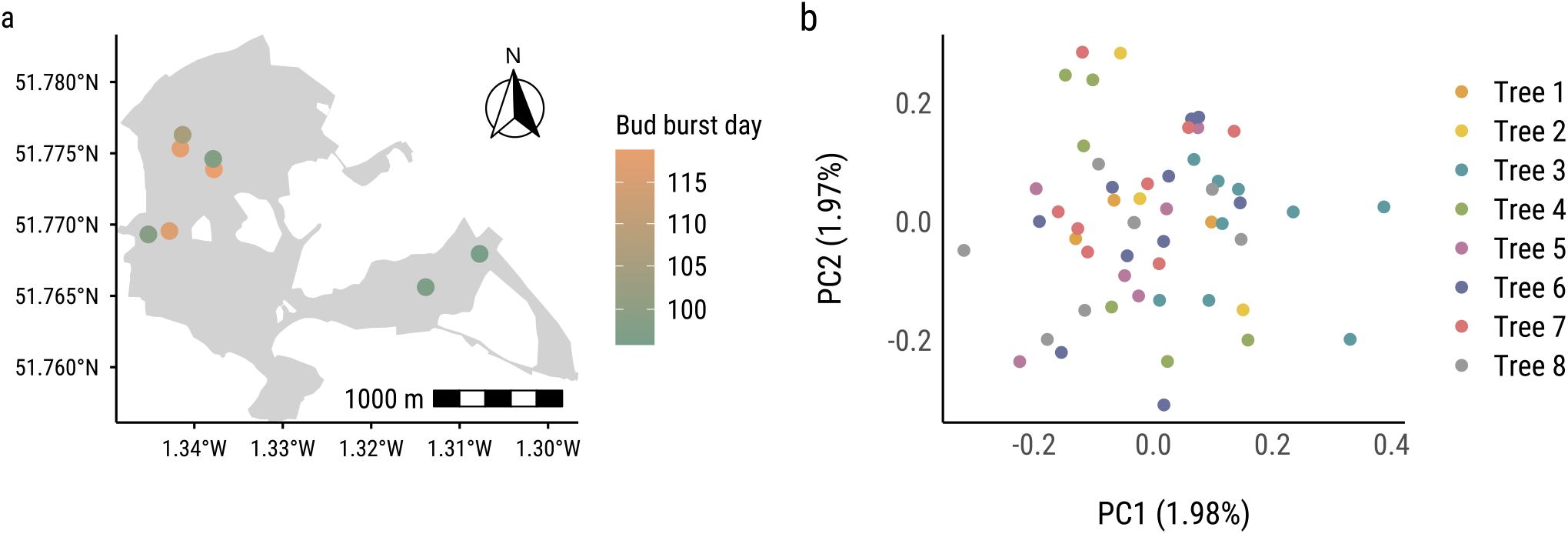
Spatial distribution of the winter moth caterpillars used for the genomic analyses and budburst timing across trees. (a) Map showing the locations of 8 individual trees within the study area, color-coded by budburst day. We sampled between 3 and 10 caterpillars per tree. (b) Genomic PCA based on autosomes reveals no clear population structure, with individuals forming a continuous distribution rather than discrete clusters associated with tree phenology.

#### DNA extraction and sequencing

We extracted DNA from the preserved caterpillars using the Qiagen DNeasy Blood & Tissue Kit, following the manufacturer’s protocol. We cleaned the samples of RNA by adding 5 µL of RNAse (10 mg*/µ*L) for 30 minutes at 37°C and removed the RNAse with the QIAamp DNA Micro Kit. We then sent the DNA extracts to Novogene UK for wholegenome sequencing using the NovaSeq X Plus platform with paired-end 150 bp reads, generating 10 GB of data per sample, providing an average of 11.5x coverage (Fig. S3).

#### Genotype calling

We trimmed adapter sequences and low-quality base calls from reads using *fastp* [45], removing the first 10 base pairs from each read with default settings and producing a quality control report. We then aligned the cleaned reads from each individual to the chromosome-level winter moth reference genome version 1.1 (ilOpeBrum1.1; `GCA_932527175.1`; [46]) using the BWA-MEM algorithm with default settings and marked and removed duplicates with Picard (http://broadinstitute.github.io/picard/). We used BCFtools [47] to call only biallelic variants, excluding those with a quality score less than or equal to 30 and a depth of coverage greater than 1500 and lower than 3, which resulted in 21,530,281 SNPs.

#### Population structure

To explore population structure, we performed a Principal Component Analysis (PCA) using PLINK [48]. We removed SNPs with a minor allele frequency (MAF) below 5%, as well as those with more than 5% missing data across individuals and removed individuals with more than 5% missing genotypes. We removed SNPs in high linkage disequilibrium (LD) because they can bias population structure inference. We removed those SNPs with an *r*^2^ higher than 0.2 in windows of 50 kb, using a step size of 50 SNPs, leaving us with 7,775,295 SNPs. We focused exclusively on autosomes. This approach helps identify whether genetic structure arises from limited gene flow due to geographical isolation or temporal isolation [49]. Including sex chromosomes could introduce bias, especially given the potential for sex-biased dispersal due to winter moth females being brachypterous.

We repeated the PCA after excluding closely related samples, which were capturing most of the variation. We estimated relatedness using the KING algorithm [50] in PLINK and removed samples related at the third degree or higher by filtering those samples with a relatedness lower than 0.059. We then plotted the eigenvectors that explained the most variance.

#### Population divergence statistics

We calculated pairwise global *F*_*ST*_ using VCFtools [46]. Populations were defined as groups of winter moths collected from the same tree. We also performed genome-wide scans by pooling individuals across trees to calculate Tajima’s D and nucleotide diversity (π) using VCFtools. To determine an appropriate window size, we examined LD decay, estimated in VCFtools as the correlation coefficient (*r*^2^) between SNP pairs. Since LD levels stabilised somewhat before 50 kb (Fig. S4), we used this window size for the analysis. This allowed us to identify any outlier regions across the population that might indicate selection or demographic processes affecting specific genomic regions.

#### Environmental associations

Although both global and windowed *F*_*ST*_ values across populations were near zero, suggesting no genetic differentiation, local adaptation could still be occurring through polygenic selection, generating subtle genomic signals that are difficult to detect using standard population differentiation metrics. To explore the potential for polygenic adaptation, we applied environmental association analyses, which associate allele frequencies with environmental variables and allow us to detect loci potentially involved in local adaptation.

We used Latent Factor Mixed Models (LFMM) as implemented in the *LEA* R package [51] to identify SNPs associated with phenological variation. LFMMs incorporate unobserved confounding variables (latent factors) to model population structure and demographic history in genomic data, thereby controlling for spurious associations that may arise from neutral evolutionary processes. Prior to LFMM analysis, we performed quality control filtering on each chromosome separately, removing SNPs with >10% missing data, monomorphic loci, and variants with MAF<0.05. We first determined the optimal number of latent factors (K) using the sparse non-negative matrix factorisation (sNMF) algorithm implemented in LEA, testing K values from 1-3 and selecting the value with the lowest cross-entropy. Based on this analysis, we used K=1 for all subsequent LFMM runs. For each chromosome, we ran LFMM using budburst date as the environmental variable of interest, with 5 repetitions, 10,000 iterations, and 5,000 burn-in steps to ensure convergence. We extracted z-scores from all repetitions and calculated the median z-score for each SNP to obtain robust test statistics. P-values were then calibrated using the genomic inflation factor (λ) calculated as the median of squared z-scores *×* divided by the median of a *χ*^2^ distribution with 1 degree of freedom. Finally, we applied a Bonferroni correction with alpha of 5 × 10^*−*5^. We validated model performance by examining p-value distributions and genomic inflation factors to ensure proper calibration of test statistics.

## Results

### Temperature manipulation experiment

Our model including temperature treatment, maternal collection day, and source tree budburst day on egg half-hatch day, with maternal identity as a group-level effect explained 97.3% of variation in half-hatch day (see Table S2 for full model summary). The mean time for a clutch to reach 50% hatched (half-hatch day) decreased significantly with increasing temperature, with a mean half-hatch day of year of 138.08 (± 0.37 SE) in the cold treatment compared to a mean of 94.60 (± 0.70 SE) in the hot treatment (Fig. 2).

Our model showed no effect of the timing of the maternal source tree at any temperature treatment (Fig. 2b), with no change in half-hatch day with each day advance or delay in budburst (estimate = -0.10, 95% CI = [-0.30, 0.10]). However half hatch day was 0.31 (95% CI = [0.17, 0.45]) days later for every day later the female was caught in the field (Fig. 2c). The effect of maternal collection day on half-hatch day was temperature-dependent, with stronger impacts at higher temperatures. Thus, the delay in egg hatching of clutches from late-emerging females was greater in the hot and warm treatments than in colder treatments.

Given that there were no significant impacts of tree phenology on egg phenology, we produced a conditional effects reaction norm at the level of the clutch to examine whether there is consistent variation in their temperature responses (Fig. 2d). The predominance of parallel lines in the reaction norm shows that clutches are strikingly consistent in their temperature response. This indicates females differ in their baseline hatching time (intercept), but show little difference in the plasticity of this trait to temperature (slope).

### Translocation experiment

Across all bags, the mean number of surviving pupae at the end was 1.01 (± 0.084 SE), from an initial number of 12 per bag, with a mean weight of 0.0323g (± 0.0008 SE). On average, bags contained 57.83 (± 1.68 SE) oak leaves with an average of 12.35 (± 0.79 SE)% of the leaf consumed, based on three sampled leaves per bag. Bags with experimentally introduced winter moth eggs experienced 6.60% (± 2.76 SE) greater herbivory than control bags (F_1,249_ = 5.71, *R*^2^ = 0.02, *p* = 0.02).

We modelled how four metrics of fitness (survival to caterpillar, survival to pupa, number of live pupae, and live pupal weight) were affected by the number of days difference in budburst relative to the parental source tree, as well as leaf number and the number of non-winter moth lepidoptera present. All model estimates for the mean weight of surviving pupae were within 0.001 of 0 (CI = [0.00, 0.00]), and were therefore excluded from Figure 3c to avoid distorting the plot. Our models showed no significant change in any fitness metric with changes in the number of days difference in budburst at the source and translocation trees (Fig. 3c). Shifts towards earlier or later trees had equal effects, as indicated by the non-significant quadratic day difference term (estimate = 0.00, 95% CI = [0.00, 0.00]). A one day shift in either direction did not affect the number of individuals surviving to caterpillar (estimate = 0.01, 95% CI = [-0.03, 0.05]), pupae (estimate = -0.01, 95% CI = [-0.06, 0.04]), or the number of live pupae (estimate = 0.02, 95% CI = [-0.03, 0.07]). Increasing numbers of non-winter moth pupae did not significantly affect the number of individuals surviving to caterpillar (estimate = 0.04, 95% CI = [-0.04, 0.12]), the number surviving to pupation (estimate = 0.02, 95% CI = [-0.11, 0.14]), or the number of live pupae (estimate = 0.10, 95% CI = [-0.02, 0.20]). Likewise the number of leaves present did not affect the number of individuals surviving to caterpillars (estimate = -0.07, 95% CI = [-0.18, 0.04]) or live pupae (estimate =, 95% CI = [-0.11, 0.29]), but was linked to slight declines in the number surviving to pupation (estimate = -0.16, 95% CI = [-0.30, -0.02]).

### Genomics

To avoid biases in the PCA caused by close familial relationships rather than true population structure, we removed one individual from each pair or group of related samples (up to the third degree). During relatedness estimation, we identified three samples with first-degree relatedness (∼0.22), all collected from the same tree using a water trap (Fig. S5). Additionally, we detected one pair of individuals with a relatedness coefficient of 0.09, consistent with either second-degree (half-sibling) or third-degree (first cousin) relationships. These two individuals were sampled from trees located approximately 1 km apart.

We conducted a PCA using autosomal SNPs to investigate potential genetic structure among individuals sampled from trees across different distances and phenologies. The first two principal components, PC1 and PC2, explained only 1.98% and 1.97% of the total genetic variance, respectively (Fig. 4b). This extremely low proportion of explained variance indicates a lack of pronounced population structure in the dataset. Additionally, individuals did not cluster by tree of origin or any other apparent grouping, suggesting high levels of genetic homogeneity across trees. We calculated global *F*_*ST*_ values for all pairwise comparisons between trees, and all estimates were 0, indicating no genetic differentiation between sampling locations.

The estimate of π was relatively uniform across all chromosomes, ranging from approximately 0.003 to 0.012, with a median of 0.0041 (Fig. 5a). This pattern suggests a lack of highly differentiated genomic regions that would create substantial heterogeneity in diversity levels. Tajima’s D values fluctuated around zero across most chromosomes, with values primarily falling within the range of -1 to +1 (Fig. 5b). Several genomic regions showed deviations from neutrality, with only two peaks on Chromosome Z exceeding the thresholds (±5 standard deviations).

**Fig. 5.**
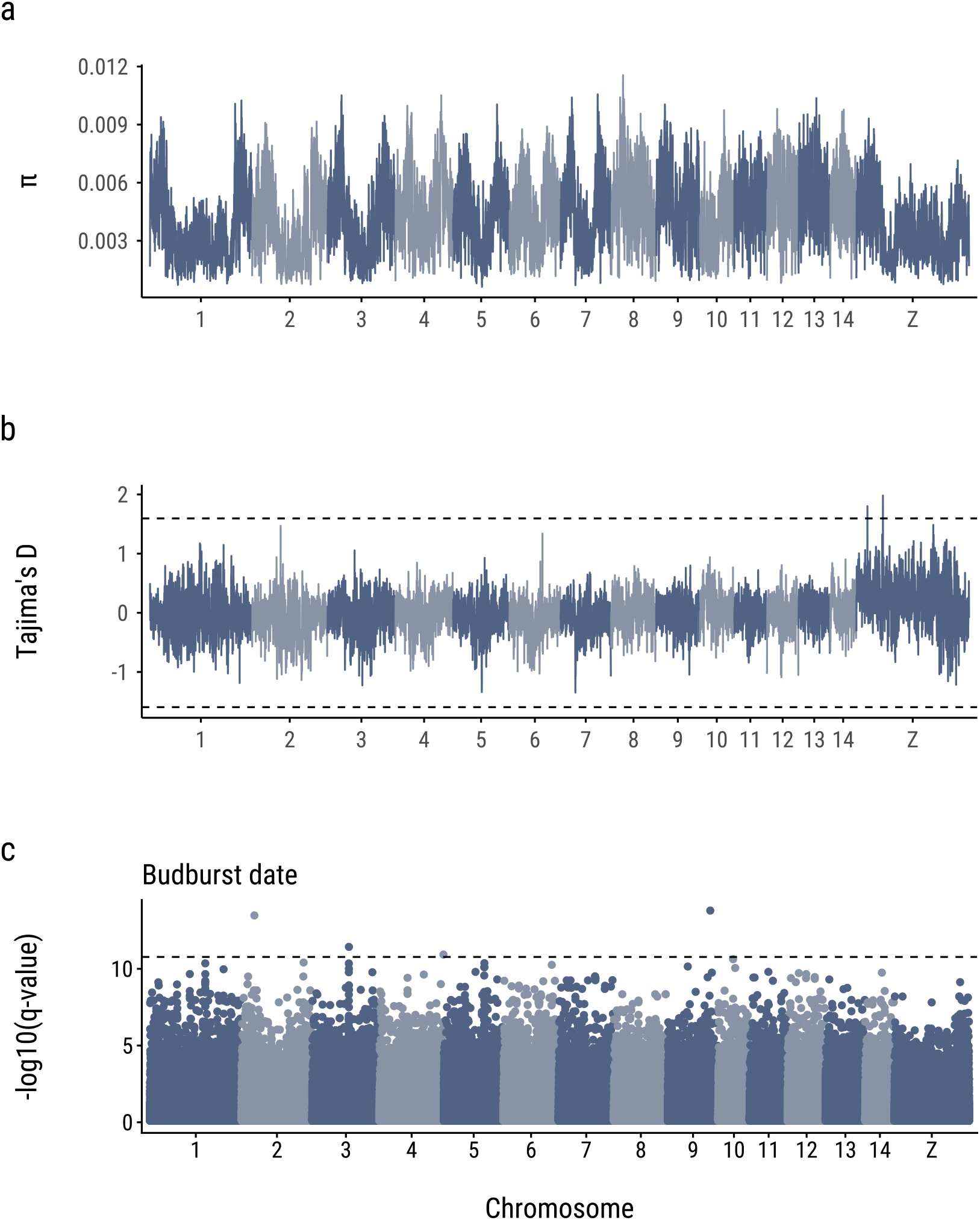
Genome-wide patterns of nucleotide diversity, population structure, and environmental association analysis. (a) Nucleotide diversity (π) across chromosomes showing relatively uniform levels of genetic diversity genome-wide. (b) Tajima’s D statistics across chromosomes with dashed lines indicating the critical thresholds (±5 standard deviations). (c) Manhattan plot resulting from an LFMM analysis of association with budburst date. Points represent SNPs with their -log(q-values) plotted against chromosomal position. The horizontal dashed line indicates the Bonferroni correction.

We applied a LFMM to identify SNPs associated with budburst date. The resulting Manhattan plot (Fig. 5c) displays only three SNPs appearing to surpass the genome-wide significance threshold with signals dispersed across multiple chromosomes. However, the corresponding QQ plot (Fig. S6) reveals that the overall distribution of test statistics is conservative, with a genomic inflation factor (λ) of 0.5. This deflation suggests that the signal-to-noise ratio is low.

## Discussion

We assessed evidence for small-scale local adaptation in winter moths to host phenology at the level of individual trees using an integrative approach, combining lab-based common garden temperature manipulations, translocation experiments in the field, and whole-genome sequencing. Our results demonstrate that despite clutch-level consistency in phenology, any small-scale selection is insufficient to overcome gene flow and produce local adaptation at this scale. Together, these findings advance our understanding of mechanisms maintaining trophic synchrony in phenologically variable woodlands.

### Temperature manipulation experiment reveals consistent relative phenology but no links to tree budburst timing

The rate of winter moth egg hatching accelerated with increasing temperatures, as has previously been reported in experimental [52,53] and long-term observational studies [54,55]. Egg hatch timing also advanced with the mother’s capture date, reflecting expected phenological carryover effects from adult eclosion timing [25,56]. We found no association between hatch timing and source tree budburst timing. This suggests that, while local adaptation to budburst timing can arise at larger scales [57], or where differences in phenology are more extreme, for example between species [58–60], the high levels of gene flow implied by our genomic results and lack of selection observed in translocation experiments do not allow local adaptation at this microgeographic scale. Based on our findings, future studies should expand the spatial scale of inquiry beyond winter moth dispersal limits where gene flow is weaker. Understanding the scales at which local adaptation in phenology can arise is a key step for understanding synchrony with local host plants across their range, and how it may drive spatial variation in phenological responses to climate change. We further report strong consistency in hatch timing at the clutch level, with clutches maintaining very similar relative timing between different temperature treatments. As egg hatch timing is a heritable trait (h^2^ = 0.63-0.94, [61,62]), this indicates different genotypes respond equally to changes in the environment, so there is no genotype-environment interaction. The uniform plastic responses across clutches suggest that selection has favoured a single, generalist developmental program that can respond appropriately to temperature cues across the range of host phenologies present within the woodland. This contrasts with scenarios where genetic adaptation would be expected to produce divergent plastic responses or reaction norm slopes tailored to specific local conditions. This plasticity-based strategy may be particularly adaptive in heterogeneous environments where gene flow prevents local genetic adaptation but environmental predictability allows plastic responses to track optimal timing. However, as winter moths tend to show greater advances in their timing with warming than host trees such as oaks [26], the lack of variation in plasticity could exacerbate mismatch with host plants, driving fitness reductions through consistent shifts towards excessively early hatching beyond the limits of buffering capacities [63].

### Translocation experiments indicate no strong selection on hatch timing synchrony

In the translocation experiment, survival and predicted fecundity were not affected by the number of days mismatched with the source tree in either direction. The lack of strong selection reported here contradicts the expectation that winter moth larvae will suffer from starvation if they hatch too early [4,10], or unpalatable, tannin-rich leaves if too late [29]. Indeed, while previous experiments have reported only 0.47-1.6% survival after hatching four to five days before budburst [4,10], clutches in this study were shifted to trees up to 13.1 days earlier than their source tree without apparent fitness consequences. The less extreme response to early hatching in our study likely arose through behavioural buffering mechanisms in the larvae. As is noted in several field guides [64,65], and has more recently been shown experimentally [66], winter moth larvae can successfully feed on unburst buds. In previous lab experiments, neonate larvae were starved until the date of budburst [4], relying solely on their starvation tolerance for survival [67]. In contrast, our field translocation placed eggs in a natural setting where they would be able to access unburst buds from the point of hatching. This suggests that often-overlooked buffering mechanisms in winter moths can compensate for mismatch and so result in weaker selection for synchrony.

### High gene flow prevents population structure

Our genomic analyses revealed extremely low genetic differentiation among winter moths collected across Wytham Woods, with the first two principal components explaining less than 2% of the total genetic variance, respectively. This minimal level of explained variance suggests virtually no population structure at the spatial scale examined, contrasting with the strong population genetic structure previously documented across broader European scales [30,31]. The complete absence of genetic differentiation (*F*_*ST*_ = 0 across all pairwise comparisons) suggests high levels of gene flow are maintained even among trees with markedly different phenological timing.

These results did not support the ADF hypothesis. It has been suggested that phenological variation could strongly drive ADF [6], as seen in galling aphid [5] and oak leafminers [68]. However, this is dependent on meeting the initial criteria for ADF, including being short-lived, sedentary specialists on a long-lived host plant [24]. Winter moths crucially diverge from these criteria in that they are not fully sedentary. The brachypterous females may disperse only a few metres, and commonly lay eggs on their source tree [69], but winged males may undergo longer range dispersal [70,71] and can transport females during copulation [65]. As well as dispersal after eclosing from pupae, winter moth larvae can disperse after hatching if food is scarce by ‘ballooning’ [70,72].

Dispersal distances via ballooning have been variously estimated as between 4.6-50 m [72] or, more recently, up to tens of kilometres with wind assistance [70,71]. Our analysis revealed an instance of second-degree relatives found approximately 1 km apart, providing direct evidence for gene flow across substantial distances within the woodland. Given the large population sizes typical of winter moths and our ability to detect this single example of distant relatedness, such long-distance dispersal events likely occur frequently enough to counteract local genetic differentiation. This level of gene flow means populations on individual trees are not isolated as demes, contradicting previous suggestions that winter moths can undergo adaptive deme formation to individual trees [25]. The high levels of nucleotide diversity observed across all chromosomes also support the interpretation of a large, panmictic population. This genetic diversity, combined with Tajima’s D values fluctuating around neutrality across most of the genome, suggests that population bottlenecks, balancing selection, or pronounced population subdivision are not major factors shaping genetic variation [73]. Instead, the uniformly high diversity levels are consistent with a large effective population size experiencing regular gene flow. The density of genetic variation we observed (approximately one SNP every 28 bp across the 619 Mb genome) is remarkably high, especially considering that the reference genome used to map the reads is from an individual belonging to the same population [46].

Our environmental association analysis using LFMM failed to identify significant genomic regions associated with bud-burst timing variation. However, the genomic inflation factor (λ = 0.5) indicates that this lack of association is not due to population structure confounding the analysis, but rather suggests that either the individual effect sizes of adaptive variants are extremely small, or that our sample size was insufficient to detect polygenic adaptation signals. The conservative distribution of test statistics implies that any genomic signatures of local adaptation to host phenology, if present, are subtle and would require substantially larger sample sizes to detect reliably. These genomic findings are consistent with our experimental findings showing no evidence for local adaptation to host tree phenology, despite the presence of heritable variation in hatch timing and clear phenological differences among host trees. Even if selection for synchrony with host phenology occurs at the level of individual trees, the continuous movement of alleles across the woodland would counteract the fixation of locally adaptive variants.

### Methodological considerations and limitations

While our combination of approaches advances our understanding of the maintenance of synchrony with phenologically variable resources at a microgeographic scale, some limitations of the methods used should be noted. Firstly, the live specimens used across the three experiments represent different families, taken from different trees within the woodland, and in different years. This spatial and temporal inconsistency means the results from each experiment cannot be directly mapped onto each other and, while we have examined multiple aspects of local adaptation, these are not done within the same samples. Secondly, though buffering may explain the weak selection on mismatch reported in the translocation experiment, it is possible the sample size of 12 eggs per bag may have been insufficient. Winter moth survival from hatching to pupation has previously been recorded as around 40% on preferred host plants under laboratory conditions [4,58,66]. Consequently, we would expect only around 4.8 larvae to survive per bag even under ideal conditions, but a difference of even one surviving pupa could represent strong selection (∼8.3%). Other sources of mortality such as predation, parasitism, and competition within the translocation bags might further obscure any patterns of selection. Given these limitations, there remains a possibility that local adaptation does occur but has not been detected, although the high level of agreement between the independent experiments suggests it is very unlikely to arise at this scale.

Broader ecological and evolutionary implications Our findings highlight important insights into plant–insect synchrony under changing environmental conditions. At fine spatial scales, we observed that phenotypic plasticity plays a stronger role than genetic adaptation in shaping winter moth responses to environmental variation. This suggests that, at this scale, populations largely track short-term changes through plasticity rather than through local adaptation. Such flexibility could buffer populations against moderate climatic fluctuations, but it can also create vulnerabilities if warming advances moth hatching phenology at a different rate than that of host plants, thereby increasing the risk of phenological mismatch.

The interplay we observed between selection and gene flow helps to clarify the evolutionary dynamics at this scale. High gene flow can constrain local adaptation, while plasticity and behavioural buffering provide resilience to current variation. However, these buffering mechanisms can come at a cost: by reducing exposure to selection, they limit the spread of beneficial alleles and the removal of maladaptive ones, thereby decreasing the population’s evolutionary potential [74]. Understanding these trade-offs is essential for predicting the long-term stability of species interactions under rapid climate change.

Finally, our results also inform applied contexts such as invasive species management. The combination of strong plasticity, behavioural flexibility, and high gene flow might underlie the winter moth’s capacity to establish and persist in novel environments. This is consistent with its history of multiple independent colonisations of North America, where it has become a major invasive defoliator of hardwoods [75]. This combination of characteristics might help winter moths stay in synchrony with new host plants and be successful in new environments: plasticity and behavioural flexibility help them adjust quickly to new conditions, and high gene flow keeps their populations diverse and increases evolutionary potential.

### Conclusions

Our integrative investigation of potential microgeographic local adaptation of winter moths to host phenology in Wytham Woods demonstrated that, despite striking consistency in relative phenology, weak selection and high levels of gene flow do not allow for local adaptation, as demonstrated by the lack of link between source tree budburst and egg hatch timing. Our findings highlight the scale-dependency of the importance of trade-offs between selection and dispersal for local adaptation and suggest that local adaptation may arise at broader scales outside of individual dispersal limits. These findings contribute to our broader understanding of how synchrony is maintained across phenologically variable landscapes and may provide insight into its resilience to future environmental change.

## Supporting information

All supplemental information

## ACKNOWLEDGEMENTS

This work was supported by grants from the UKRI Frontiers award EP/X024520/1 to BCS. We thank Sam Crofts and Lucy Morley for help with field collection, and Natalie van Dis for discussions.

## Author contributions

Conceptualisation: RL, AE, EFC, BCS; Data collection: RL, LB; Lab work: AE (molecular), LB (molecular, experimental), RL (experimental); Formal Analysis: RL (Experiments), AE (Genomics); Funding Acquisition: BCS; Investigation: RL, AE; Methodology: RL, AE; Visualization: RL, AE; Writing - Original Draft: RL, AE; Writing - Review Editing: RL, AE, LB, EFC, BCS.

## Code availability

Code to reproduce all analyses conducted in this paper can be found at https://github.com/andreaestandia/winter-mothwytham.

## References

1. Ponti R, Sannolo M. 2023 The importance of including phenology when modelling species ecological niche. Ecography 2023, e06143. (doi:10.1111/ecog.06143)

2. Chmura HE, Kharouba HM, Ashander J, Ehlman SM, Rivest EB, Yang LH. 2019 The mechanisms of phenology: the patterns and processes of phenological shifts. Ecol. Monogr. 89, e01337. (doi:10.1002/ecm.1337)

3. Parmesan C, Yohe G. 2003 A globally coherent fingerprint of climate change impacts across natural systems. Nature 421, 37–42. (doi:10.1038/nature01286)

4. van Dis NE, Sieperda G-J, Bansal V, van Lith B, Wertheim B, Visser ME. 2023 Phenological mismatch affects individual fitness and population growth in the winter moth. Proc. R. Soc. B Biol. Sci. 290, 20230414. (doi:10.1098/rspb.2023.0414)

5. Komatsu T, Akimoto S. 1995 Genetic differentiation as a result of adaptation to the phenologies of individual host trees in the galling aphid Kaltenbachiella japonica. Ecol. Entomol. 20, 33–42. (doi:10.1111/j.1365-2311.1995.tb00426.x)

6. Mopper S. 2005 Phenology — how time creates spatial structure in endophagous insect populations. Ann. Zool. Fenn. 42, 327–333.

7. Posledovich D, Toftegaard T, Wiklund C, Ehrlén J, Gotthard K. 2015 The developmental race between maturing host plants and their butterfly herbivore – the influence of phenological matching and temperature. J. Anim. Ecol. 84, 1690–1699. (doi:10.1111/1365-2656.12417)

8. Rauschkolb R et al. 2024 Spatial variability in herbaceous plant phenology is mostly explained by variability in temperature but also by photoperiod and functional traits. Int. J. Biometeorol. 68, 761–775. (doi:10.1007/s00484-024-02621-9)

9. Cole EF, Sheldon BC. 2017 The shifting phenological landscape: Within- and between-species variation in leaf emergence in a mixed-deciduous woodland. Ecol. Evol. 7, 1135–1147. (doi:10.1002/ece3.2718)

10. van Asch M, Julkunen-Tiito R, Visser ME. 2010 Maternal effects in an insect herbivore as a mechanism to adapt to host plant phenology. Funct. Ecol. 24, 1103–1109. (doi:10.1111/j.1365-2435.2010.01734.x)

11. Phillimore AB, Stålhandske S, Smithers RJ, Bernard R. 2012 Dissecting the Contributions of Plasticity and Local Adaptation to the Phenology of a Butterfly and Its Host Plants. Am. Nat. 180, 655–670. (doi:10.1086/667893)

12. Fox RJ, Donelson JM, Schunter C, Ravasi T, Gaitán-Espitia JD. 2019 Beyond buying time: the role of plasticity in phenotypic adaptation to rapid environmental change. Philos. Trans. R. Soc. B Biol. Sci. 374, 20180174. (doi:10.1098/rstb.2018.0174)

13. Berlocher SH, Feder JL. 2002 Sympatric Speciation in Phytophagous Insects: Moving Beyond Controversy? Annu. Rev. Entomol. 47, 773–815. (doi:10.1146/annurev.ento.47.091201.145312)

14. Helm B, Doren BMV, Hoffmann D, Hoffmann U. 2019 Evolutionary Response to Climate Change in Migratory Pied Flycatchers. Curr. Biol. 29, 3714–3719.e4. (doi:10.1016/j.cub.2019.08.072)

15. Visser ME, Both C. 2005 Shifts in phenology due to global climate change: the need for a yardstick. Proc. R. Soc. B Biol. Sci. 272, 2561–2569. (doi:10.1098/rspb.2005.3356)

16. Forrest JR. 2016 Complex responses of insect phenology to climate change. Curr. Opin. Insect Sci. 17, 49–54. (doi:10.1016/j.cois.2016.07.002)

17. Thackeray SJ et al. 2010 Trophic level asynchrony in rates of phenological change for marine, freshwater and terrestrial environments. Glob. Change Biol. 16, 3304–3313. (doi:10.1111/j.1365-2486.2010.02165.x)

18. Hodgson JA, Thomas CD, Oliver TH, Anderson BJ, Brereton TM, Crone EE. 2011 Predicting insect phenology across space and time. Glob. Change Biol. 17, 1289–1300. (doi:10.1111/j.1365-2486.2010.02308.x)

19. Cogni R, Futuyma DJ. 2009 Local adaptation in a plant herbivore interaction depends on the spatial scale. Biol. J. Linn. Soc. 97, 494–502. (doi:10.1111/j.1095-8312.2009.01234.x)

20. Dittmar EL, Schemske DW. 2023 Temporal Variation in Selection Influences Microgeographic Local Adaptation. Am. Nat. 202, 471–485. (doi:10.1086/725865)

21. von Takach B, Ahrens CW, Lindenmayer DB, Banks SC. 2021 Scale-dependent signatures of local adaptation in a foundation tree species. Mol. Ecol. 30, 2248–2261. (doi:10.1111/mec.15894)

22. Blanquart F, Kaltz O, Nuismer SL, Gandon S. 2013 A practical guide to measuring local adaptation. Ecol. Lett. 16, 1195–1205. (doi:10.1111/ele.12150)

23. Edmunds GF, Alstad DN. 1978 Coevolution in Insect Herbivores and Conifers. Science 199, 941–945. (doi:10.1126/science.199.4332.941)

24. Zandt PAV, Mopper S. 1998 A Meta-Analysis of Adaptive Deme Formation in Phytophagous Insect Populations. Am. Nat. 152, 595–604. (doi:10.1086/286192)

25. Van Dongen S, Backeljau T, Matthysen E, Dhondt AA. 1997 Synchronization of Hatching Date with Budburst of Individual Host Trees (Quercus robur) in the Winter Moth (Operophtera brumata) and its Fitness Consequences. J. Anim. Ecol. 66, 113–121. (doi:10.2307/5969)

26. van Asch M, Salis L, Holleman LJM, van Lith B, Visser ME. 2013 Evolutionary response of the egg hatching date of a herbivorous insect under climate change. Nat. Clim. Change 3, 244–248. (doi:10.1038/nclimate1717)

27. Dis NE van. 2024 Genetic adaptation to climate change in wild populations: a systematic literature review identifies opportunities to strengthen our evidence base.

28. Feeny P. 1970 Seasonal Changes in Oak Leaf Tannins and Nutrients as a Cause of Spring Feeding by Winter Moth Caterpillars. Ecology 51, 565–581. (doi:10.2307/1934037)

29. Feeny PP. 1968 Effect of oak leaf tannins on larval growth of the winter moth Operophtera brumata. J. Insect Physiol. 14, 805–817. (doi:10.1016/0022-1910(68)90191-1)

30. Andersen JC, Havill NP, Mannai Y, Ezzine O, Dhahri S, Ben Jamâa ML, Caccone A, Elkinton JS. 2019 Identification of winter moth (Operophtera brumata) refugia in North Africa and the Italian Peninsula during the last glacial maximum. Ecol. Evol. 9, 13931–13941. (doi:10.1002/ece3.5830)

31. Andersen JC, Havill NP, Caccone A, Elkinton JS. 2017 Postglacial recolonization shaped the genetic diversity of the winter moth (Operophtera brumata) in Europe. Ecol. Evol. 7, 3312–3323. (doi:10.1002/ece3.2860)

32. Varley GC, Gradwell GR. 1968 Population Models for the Winter Moth. In Symposium of the Royal entomological Society of London, pp. 132–142. T. R. E. Southwood.

33. Lack D. 1964 A Long-Term Study of the Great Tit (Parus major). J. Anim. Ecol. 33, 159–173. (doi:10.2307/2437)

34. Jaworski T, Sukovata L. 2020 Tools for monitoring oak defoliating geometrids – traps for catching males and females. Scand. J. For. Res. 35, 506–512. (doi:10.1080/02827581.2020.1839125)

35. Morley LM, Crofts SJ, Cole EF, Sheldon BC. 2025 Quantifying Phenology in the Deciduous Tree and Phytophagous Insect System: A Methodological Comparison. Ecol. Evol. 15, e71821. (doi:10.1002/ece3.71821)

36. Klosterman S, Richardson AD. 2017 Observing Spring and Fall Phenology in a Deciduous Forest with Aerial Drone Imagery. Sensors 17, 2852. (doi:10.3390/s17122852)

37. Hinks AE, Cole EF, Daniels KJ, Wilkin TA, Nakagawa S, Sheldon BC. 2015 Scale-Dependent Phenological Synchrony between Songbirds and Their Caterpillar Food Source. Am. Nat. 186, 84–97. (doi:10.1086/681572)

38. Met Office, Hollis D, McCarthy M, Kendon M, Legg T. 2022 HadUK-Grid Gridded Climate Observations on a 25km grid over the UK, v1.1.0.0 (1836-2021).

39. Luedeling E, Caspersen L, Fernández E. 2024 chillR: Statistical Methods for Phenology Analysis in Temperate Fruit Trees.

40. R Core Team. 2013 R: A language and environment for statistical computing.

41. Learmonth R. Unpublished data.

42. Bürkner P-C. 2017 brms: An R Package for Bayesian Multilevel Models Using Stan. J. Stat. Softw. 80, 1–28. (doi:10.18637/jss.v080.i01)

43. Getman-Pickering ZL, Campbell A, Aflitto N, Grele A, Davis JK, Ugine TA. 2020 LeafByte: A mobile application that measures leaf area and herbivory quickly and accurately. Methods Ecol. Evol. 11, 215–221. (doi:10.1111/2041-210X.13340)

44. Kawecki TJ, Ebert D. 2004 Conceptual issues in local adaptation. Ecol. Lett. 7, 1225–1241. (doi:10.1111/j.1461-0248.2004.00684.x)

45. Chen S, Zhou Y, Chen Y, Gu J. 2018 fastp: an ultra-fast all-in-one FASTQ preprocessor. Bioinformatics 34, i884–i890. (doi:10.1093/bioinformatics/bty560)

46. Crowley LM, Sheldon BC, University of Oxford and Wytham Woods Genome Acquisition Lab, Darwin Tree of Life Barcoding collective, Wellcome Sanger Institute Tree of Life programme, Wellcome Sanger Institute Scientific Operations: DNA Pipelines collective, Tree of Life Core Informatics collective, Darwin Tree of Life Consortium. 2023 The genome sequence of the Winter Moth, Operophtera brumata (Linnaeus, 1758). Wellcome Open Res. 8, 530. (doi:10.12688/wellcomeopenres.20361.1)

47. Danecek P et al. 2011 The variant call format and VCFtools. Bioinformatics 27, 2156–2158. (doi:10.1093/bioinformatics/btr330)

48. Purcell S et al. 2007 PLINK: A Tool Set for Whole-Genome Association and Population-Based Linkage Analyses. Am. J. Hum. Genet. 81, 559–575. (doi:10.1086/519795)

49. Novembre J et al. 2008 Genes mirror geography within Europe. Nature 456, 98–101. (doi:10.1038/nature07331)

50. Manichaikul A, Mychaleckyj JC, Rich SS, Daly K, Sale M, Chen W-M. 2010 Robust relationship inference in genome-wide association studies. Bioinformatics 26, 2867–2873. (doi:10.1093/bioinformatics/btq559)

51. Frichot E, Schoville SD, Bouchard G, François O. 2013 Testing for associations between loci and environmental gradients using latent factor mixed models. Mol. Biol. Evol. 30, 1687–1699. (doi:10.1093/molbev/mst063)

52. Buse A, Good J. 1996 Synchronization of larval emergence in winter moth (Operophtera brumata L.) and budburst in pedunculate oak (Quercus robur L.) under simulated climate change. Ecol. Entomol. 21, 335–343. (doi:10.1046/j.1365-2311.1996.t01-1-00001.x)

53. Kimberling DN, Miller JC. 1988 Effects of temperature on larval eclosion of the winter moth, Operophtera brumata. Entomol. Exp. Appl. 47, 249–254. (doi:10.1111/j.1570-7458.1988.tb01143.x)

54. Salis L, Lof M, Van Asch M, Visser ME. 2016 Modeling winter moth Operophtera brumata egg phenology: nonlinear effects of temperature and developmental stage on developmental rate. Oikos 125, 1772–1781. (doi:10.1111/oik.03257)

55. Visser ME, Holleman LJM. 2001 Warmer springs disrupt the synchrony of oak and winter moth phenology. Proc. R. Soc. Lond. B Biol. Sci. 268, 289–294. (doi:10.1098/rspb.2000.1363)

56. Salis L, van den Hoorn E, Beersma DGM, Hut RA, Visser ME. 2018 Photoperiodic cues regulate phenological carry-over effects in an herbivorous insect. Funct. Ecol. 32, 171–180. (doi:10.1111/1365-2435.12953)

57. Speyer W. 1941 Beobachtungen über das Zahlenverhältnis der Geschlechter bei der Kirschfruchtfliege (Rhagoletis cerasi L.). Z. Für Pflanzenkrankh. Pflanzenpathol. Pflanzenschutz 51, 330–332.

58. Kerslake JE, Hartley SE. 1997 Phenology of Winter Moth Feeding on Common Heather: Effects of Source Population and Experimental Manipulation of Hatch Dates. J. Anim. Ecol. 66, 375–385. (doi:10.2307/5983)

59. Tikkanen O-P, Woodcock B, Watt A, Lock K. 2006 Are polyphagous geometrid moths with flightless females adapted to budburst phenology of local host species? Oikos 112, 83–90. (doi:10.1111/j.0030-1299.2006.13855.x)

60. Molleman F, Walczak U. 2024 Winter moth populations are isolated on co-occurring tree species with contrasting budburst-phenology. Ecol. Entomol. 49, 744–747. (doi:10.1111/een.13351)

61. Tikkanen O-P, Lyytikäinen-Saarenmaa P. 2002 Adaptation of a generalist moth, Operophtera brumata, to variable budburst phenology of host plants. Entomol. Exp. Appl. 103, 123–133. (doi:10.1046/j.1570-7458.2002.00966.x)

62. Van Asch M, Van Tienderen PH, Holleman LJM, Visser ME. 2007 Predicting adaptation of phenology in response to climate change, an insect herbivore example. Glob. Change Biol. 13, 1596–1604. (doi:10.1111/j.1365-2486.2007.01400.x)

63. Weir JC. 2024 Trophic generalism in the winter moth: a model species for phenological mismatch. Oecologia 206, 225–239. (doi:10.1007/s00442-024-05629-5)

64. Newman E. 1870 An Illustrated Natural History of British Butterflies and Moths. Hardwicke and Bogue.

65. Stokoe WJ. 1948 The caterpillar of British moths. London: Warne.

66. Weir JC. 2023 Buffering and trophic mismatch in spring-feeding forest caterpillars. (doi:10.7488/era/3194)

67. Tikkanen O-P, Julkunen-Tiitto R. 2003 Phenological variation as protection against defoliating insects: the case of Quercus robur and Operophtera brumata. Oecologia 136, 244–251. (doi:10.1007/s00442-003-1267-7)

68. Mopper S, Beck M, Simberloff D, Stiling P. 1995 Local Adaptation and Agents of Selection in a Mobile Insect. Evolution 49, 810–815. (doi:10.1111/j.1558-5646.1995.tb02317.x)

69. Graf B, Borer F, Höpli HU, Höhn H, Dorn S. 1995 The winter moth, Operophtera brumata L. (Lep., Geometridae), on apple and cherry: spatial and temporal aspects of recolonization in autumn. J. Appl. Entomol. 119, 295–301. (doi:10.1111/j.1439-0418.1995.tb01289.x)

70. Vindstad OPL, Jepsen JU, Yoccoz NG, Bjørnstad ON, Mesquita M d. S, Ims RA. 2019 Spatial synchrony in sub-arctic geometrid moth outbreaks reflects dispersal in larval and adult life cycle stages. J. Anim. Ecol. 88, 1134–1145. (doi:10.1111/1365-2656.12959)

71. Vindstad OPL, Jepsen JU, Molvig H, Ims RA. 2022 A pioneering pest: the winter moth (Operophtera brumata) is expanding its outbreak range into Low Arctic shrub tundra. Arct. Sci. 8, 450–470. (doi:10.1139/as-2021-0027)

72. Edland T. 1971 Wind dispersal of the winter moth Operophtera brumata L. (Lep., Geometridae) and its relevance to control measures. Nor. Entomol. Tidskr. 18, 103–105.

73. Nielsen R. 2001 Statistical tests of selective neutrality in the age of genomics. Heredity 86, 641–647. (doi:10.1046/j.1365-2540.2001.00895.x)

74. Rice KJ, Emery NC. 2003 Managing microevolution: restoration in the face of global change. Front. Ecol. Environ. 1, 469–478. (doi:10.1890/1540-9295(2003)001[0469:MMRITF]2.0.CO;2)

75. Andersen JC, Havill NP, Caccone A, Elkinton JS. 2021 Four times out of Europe: Serial invasions of the winter moth, Operophtera brumata, to North America. Mol. Ecol. 30, 3439–3452. (doi:10.1111/mec.15983)

