## Supplementary material for "Integrative analysis of fine-scale local adaptation of winter moths to variable oak phenology": All supplemental information

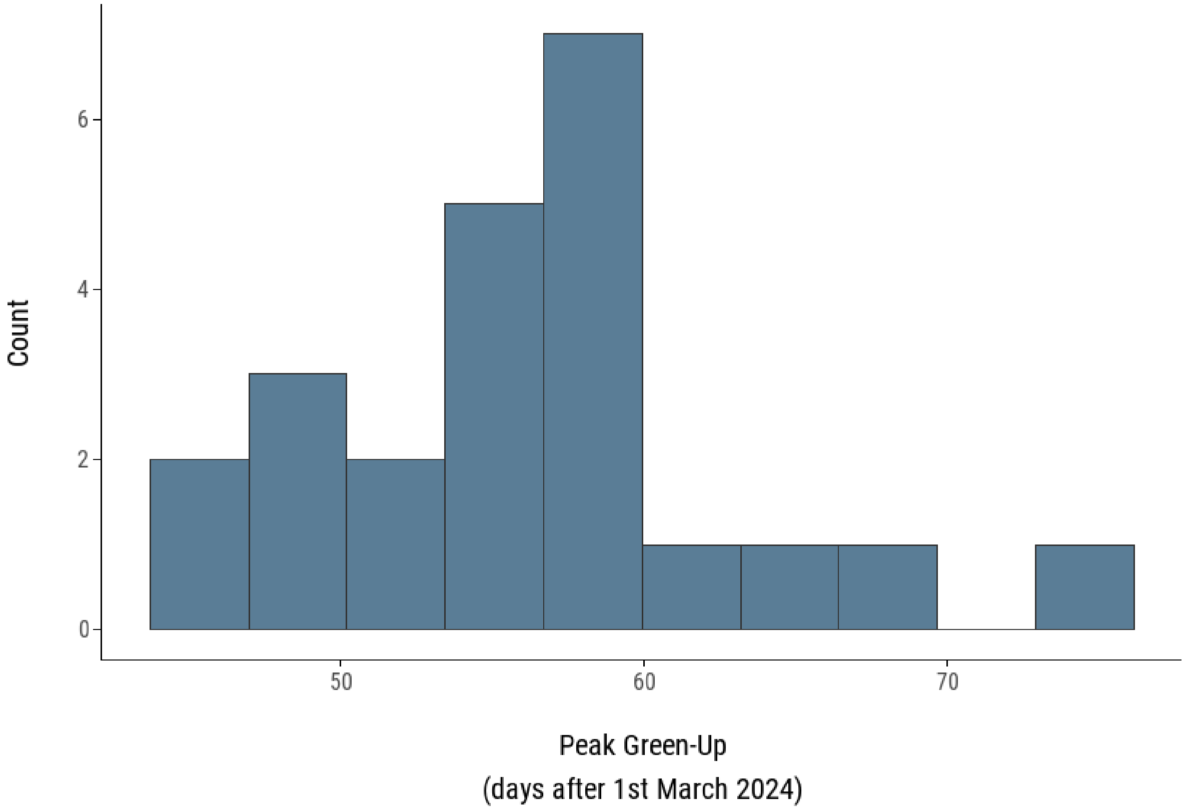


***SI***

***SI Figure 1: Histogram of 2024 green-up dates for Wytham transect trees where adult moths were sampled in 2024****. The timing of peak green-up covers a 29-day range and conforms to a normal distribution, based on a Shapiro-Wilk normality test (W = 0.94, p-value = 0.29).*

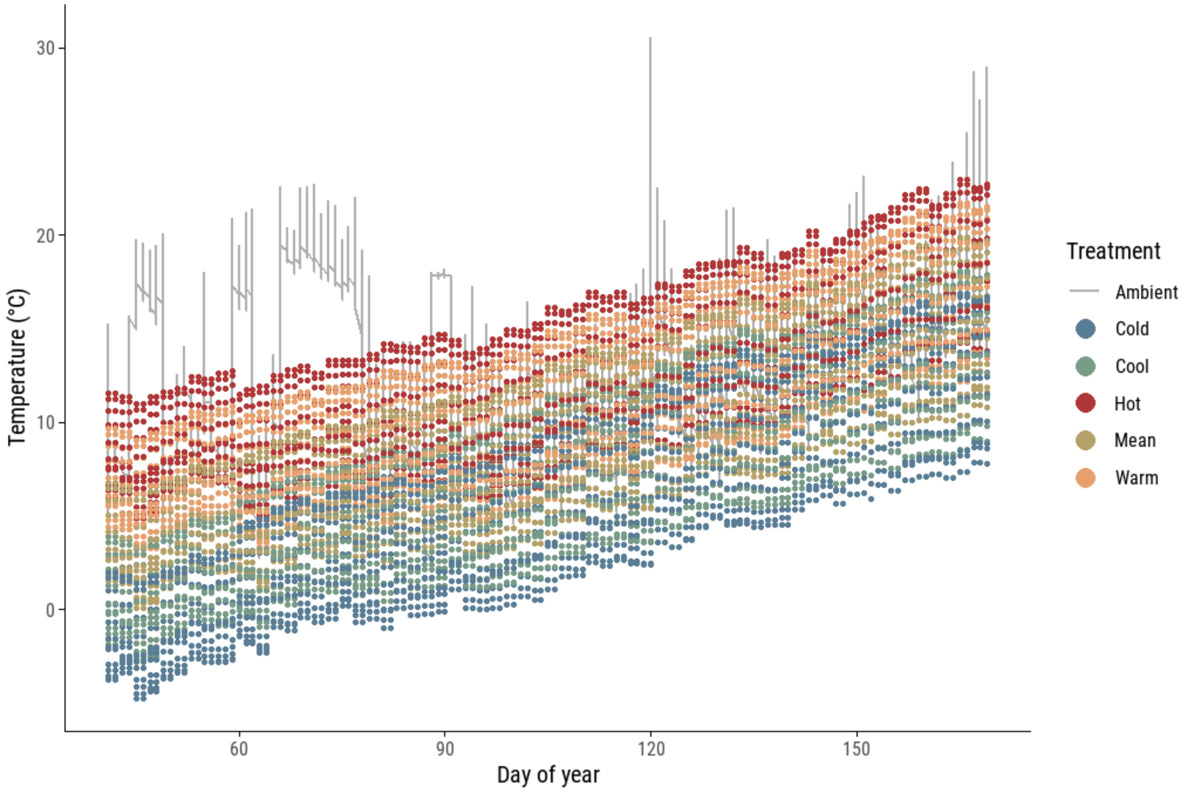


***SI Figure 2: Temperature treatments across five incubators, and ambient outdoor treatment plotted over time for 2025 up to timing of final egg hatching in the cold treatment.*** *Each experimental treatment is plotted as dots in the colours shown in the key, while the ambient outdoor treatment is plotted as a grey line.*

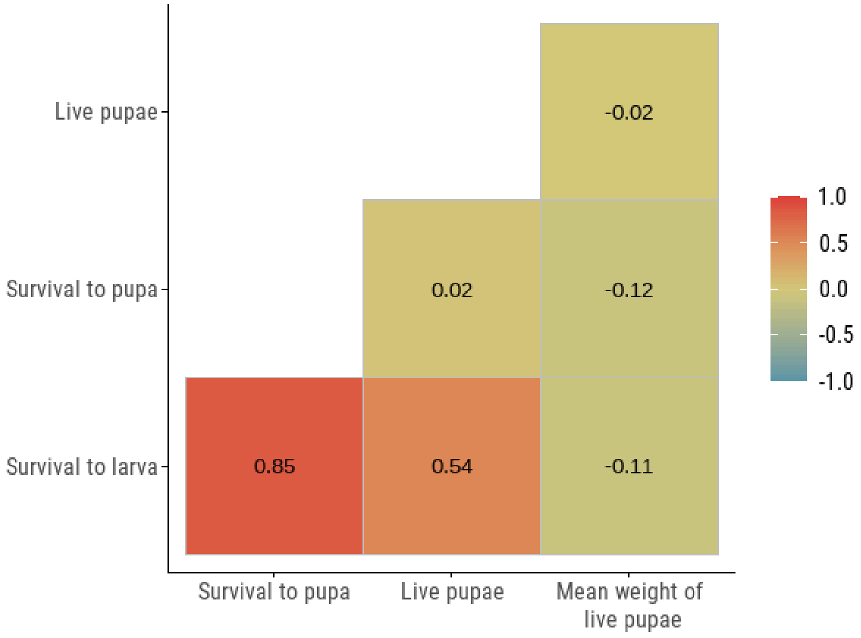


***SI Figure 3: Pearson correlation matrix of the four fitness metrics considered in translocation experiments.*** *The colour indicates the strength of the correlation, while the number indicates the Pearson correlation coefficient.*

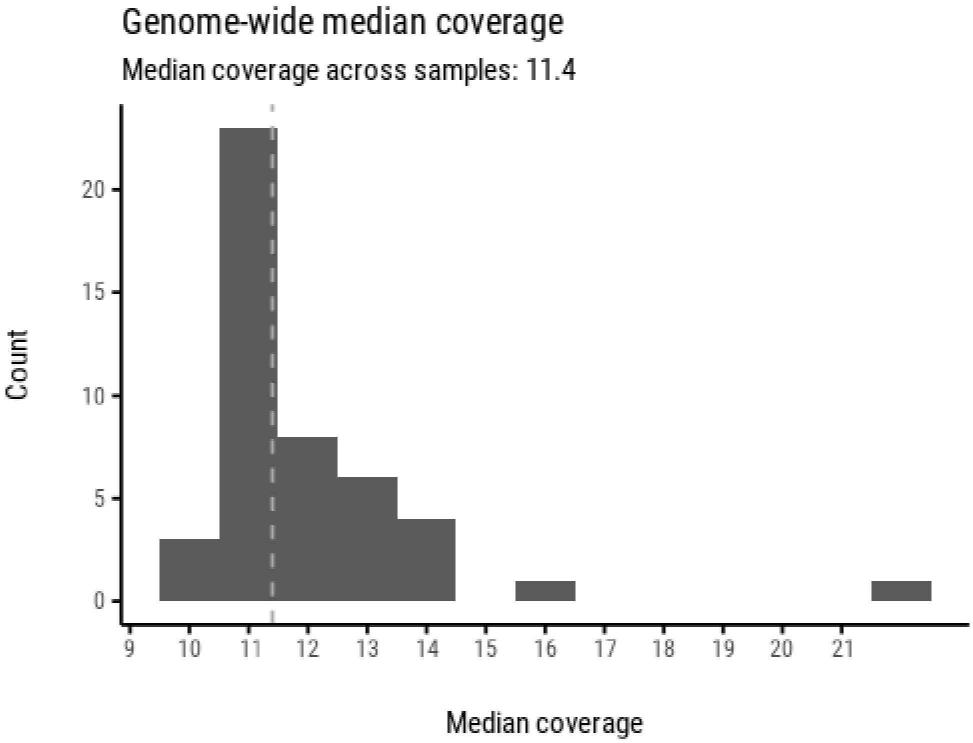


***SI Figure 4*. *Distribution of genome-wide median sequencing coverage across samples.*** *Histogram bars show the number of samples (y-axis) falling into each median coverage bin, with a dashed vertical line indicating the overall median coverage of 11.4×.*

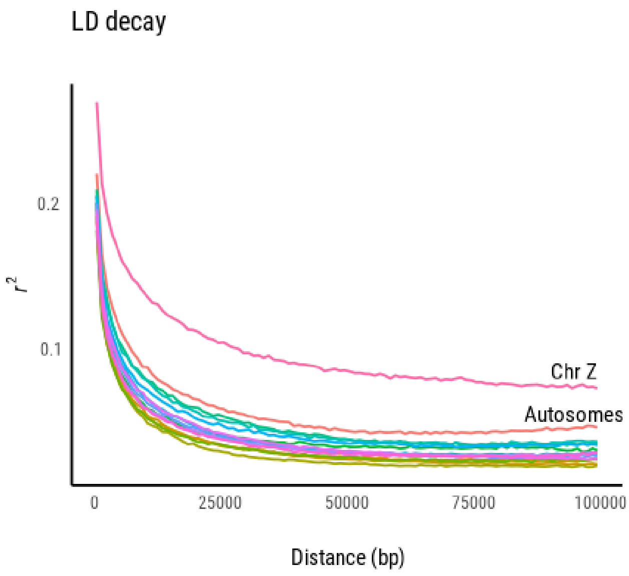


***SI Figure 5. Linkage disequilibrium (LD) decay across chromosomes.*** *The plot shows the average pairwise LD (measured as $r^2$) as a function of physical distance (bp) for different chromosomes. The Z chromosome displays higher LD over longer distances compared to autosomes, consistent with reduced effective recombination and sex-specific inheritance patterns. LD decays more rapidly within the first 50 kb and stabilises beyond that distance.*

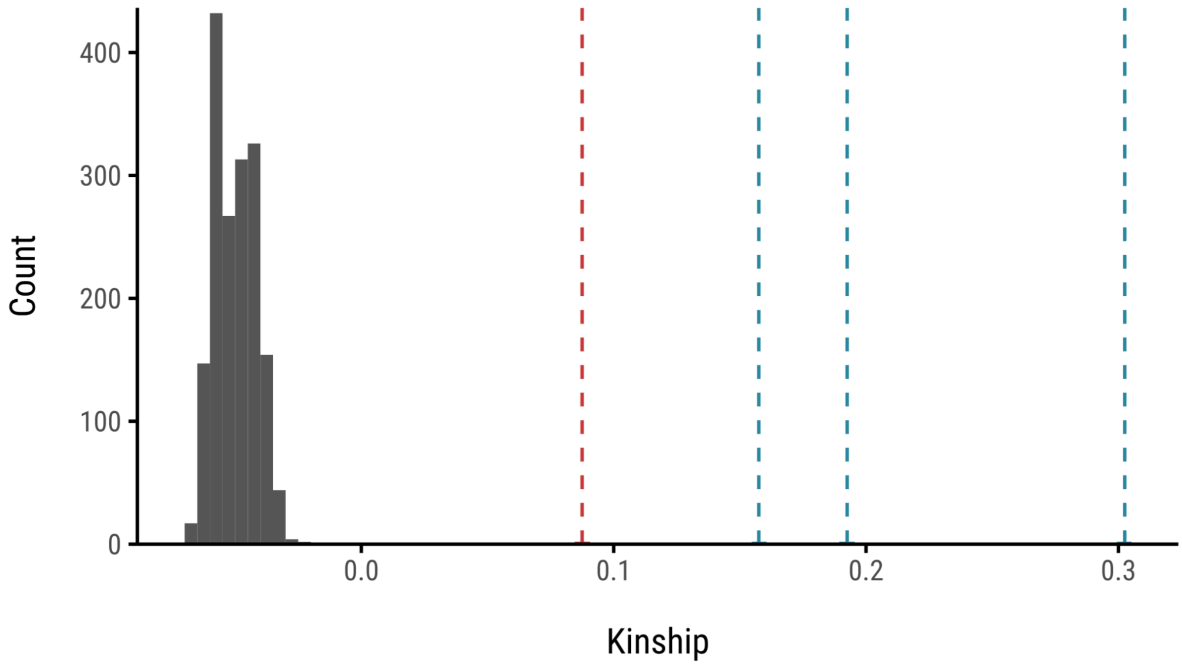


***SI Figure 6*. *Distribution of kinship coefficients calculated using KING.*** *The red vertical line marks a pair of second-degree relatives (e.g., half-siblings) sampled from two trees separated by ~1 km. The blue vertical lines indicate three individuals with first-degree relationships, all collected from the same tree using a water trap, likely representing full siblings. Note that negative values are possible and they typically indicate unrelated relationships or ancestry differences.*

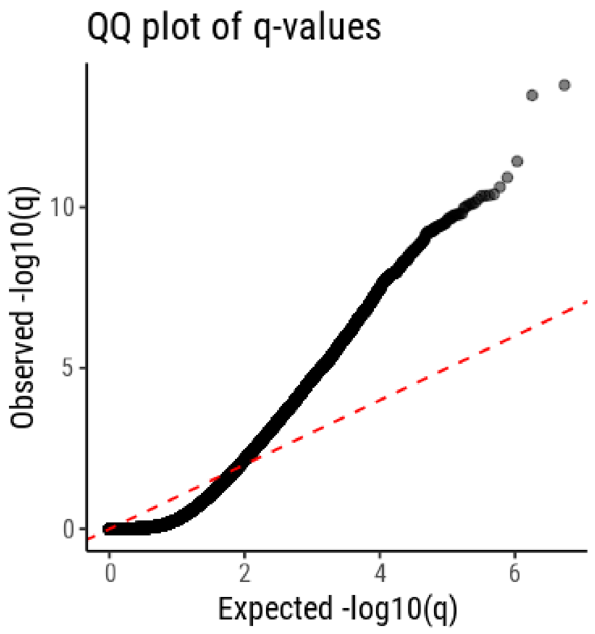


***SI Figure 7***. *QQ plot comparing observed and expected –$log_{10}$(q-values) from the LFMM analysis. The genomic inflation factor (α = 0.5) indicates conservative test statistics, with most points falling below the red dashed line representing the null expectation.*

***SI Table 1: Distribution of origin of clutches in temperature manipulation experiment.*** *The table lists the trees from which female winter moths were sampled, their 2025 half-hatch date as determined by the half-point of maximum budburst score, from ground observations, and the number of clutches used in the temperature manipulation experiment originating from each tree.*

| Mother's source tree | Half-date of source tree | Number of clutches in experiment |
|---|---|---|
| 14128 | NA | 8 |
| 17321 | NA | 2 |
| 20708 | NA | 1 |
| T11 | 98.65836 | 4 |
| T12 | 94.33183 | 5 |
| T13 | 98.2352 | 7 |
| T14 | 98.39315 | 10 |
| T15 | 99.52082 | 1 |
| T16 | 98.35866 | 1 |
| T17 | 109.2942 | 1 |
| T18 | 117.7242 | 1 |
| T19 | 102.7511 | 3 |
| T2 | 102.2977 | 1 |
| T20 | 104.0373 | 5 |
| T3 | 100.531 | 10 |
| T4 | 104.2981 | 9 |
| T5 | 96.04565 | 1 |
| T6 | 100.7654 | 1 |
| T7 | 103.0496 | 1 |
| T8 | 96.7073 | 2 |
| T9 | 100.2441 | 1 |

**SI Table 2:** ***Summary of population-level effects for modelling the determinants of half-hatch date in the temperature manipulation experiment.*** *The estimate, estimate error, upper and lower 95% confidence interval limits, Rhat, bulk, and tail effective sample sizes (ESS) are given. Rhat represents the potential scale reduction factor for the four chains and represents convergence at 1 for all variables.*

| | Estimate | Est.Error | Lower 95% CI | Upper 95% CI | Rhat | Bulk ESS | Tail ESS |
|---|---|---|---|---|---|---|---|
| **Intercept** | 25.16 | 28.64 | -31.90 | 81.13 | 1.00 | 3103 | 5098 |
| **Cold Treatment** | 77.39 | 23.43 | 31.64 | 122.51 | 1.00 | 3785 | 5960 |
| **Cool Treatment** | 59.40 | 23.49 | 13.34 | 104.50 | 1.00 | 3696 | 5967 |
| **Warm Treatment** | -32.35 | 23.72 | -79.5 | 14.21 | 1.00 | 3625 | 5151 |
| **Hot treatment** | -59.60 | 23.59 | -105.22 | -13.29 | 1.00 | 3909 | 5472 |
| **Ambient treatment** | 3.22 | 23.65 | -43.82 | 49.29 | 1.00 | 3527 | 5336 |
| **Maternal collection day** | 0.31 | 0.07 | 0.17 | 0.45 | 1.00 | 2970 | 4990 |
| **2025 source tree budburst day** | -0.10 | 0.10 | -0.30 | 0.10 | 1.00 | 3992 | 5557 |
| **Cold: maternal collection day** | -0.17 | 0.07 | -0.31 | -0.04 | 1.00 | 3801 | 5945 |
| **Cool: maternal collection day** | -0.14 | 0.07 | -0.27 | -0.01 | 1.00 | 3709 | 6059 |
| **Warm: maternal collection day** | 0.05 | 0.07 | -0.09 | 0.19 | 1.00 | 3634 | 5031 |
| **Hot: maternal collection day** | 0.10 | 0.07 | -0.03 | 0.24 | 1.00 | 3917 | 5367 |
| **Ambient: maternal collection day** | -0.06 | 0.07 | -0.20 | 0.08 | 1.00 | 3533 | 5427 |
